# First whole genome assembly of *Vaccinium floribundum* Kunth, an emblematic Andean species

**DOI:** 10.1101/2024.04.26.591377

**Authors:** Martina Albuja-Quintana, Gabriela Pozo, Milton Gordillo-Romero, Carolina E. Armijos, María de Lourdes Torres

## Abstract

**Background:** *Vaccinium floribundum* Kunth, known as "mortiño," is an endemic shrub species of the Andean region adapted to harsh conditions in high-altitude ecosystems. It plays an important ecological role as a pioneer species in the aftermath of deforestation and human-induced fires within paramo ecosystems, emphasizing its conservation value. While previous studies have offered insights into the genetic diversity of mortiño, comprehensive genomic studies are still missing to fully understand the unique adaptations of this species and its population status, highlighting the importance of generating a reference genome for this plant.

**Results:** ONT and Illumina sequencing were used to establish a reference genome for this species. Three different *de novo* genome assemblies were generated and compared for quality, continuity and completeness. The Flye assembly was selected as the best and refined by filtering out short ONT reads, screening for contaminants and genome scaffolding. The final assembly has a genome size of 529 MB, containing 1,317 contigs and 97% complete BUSCOs, indicating a high level of integrity of the genome. Additionally, the LAI Index of 12.93, further categorizes this assembly as a reference genome.

**Conclusions:** The genome of *V. floribundum* reported in this study is the first reference genome generated for this species, providing a valuable tool for further studies. This high-quality genome, based on the quality and completeness parameters obtained, will not only help uncover the genetic mechanisms responsible for its unique traits and adaptations to high-altitude ecosystems, but will also contribute to conservation strategies for a species endemic to the Andes.

## Introduction

The Andean blueberry (*Vaccinium floribundum* Kunth), also known as mortiño (**Figure 1**), is a diploid (2n=2x=24) wild perennial shrub species native to the Andean region, including the highlands of Venezuela, Colombia, Bolivia, Perú and Ecuador [1–5]. In Ecuador, *V. floribundum* is recognized as a highly valued wild crop with economic and cultural value owing to the nutritional richness and exquisite flavor of its berries. These berries are traditionally used in the preparation of wine, ice creams, and marmalade, and they also serve as an essential ingredient in the traditional “colada morada” beverage consumed during the Day of the Souls (“Día de los Muertos”) in Ecuador [6–8]. Furthermore, the fruits of *V. floribundum* boast a remarkable nutritional profile, attributed to the presence of phytochemical nutrients with antioxidant capacity, including polyphenols, anthocyanins and flavonols [9–11]. Additionally, these fruits harbor an array of bioactive compounds that confer multiple health benefits, including anti-inflammatory [12], antibacterial [13], antitumor [14] and antihypertensive properties [4,15].

**Figure 1.**
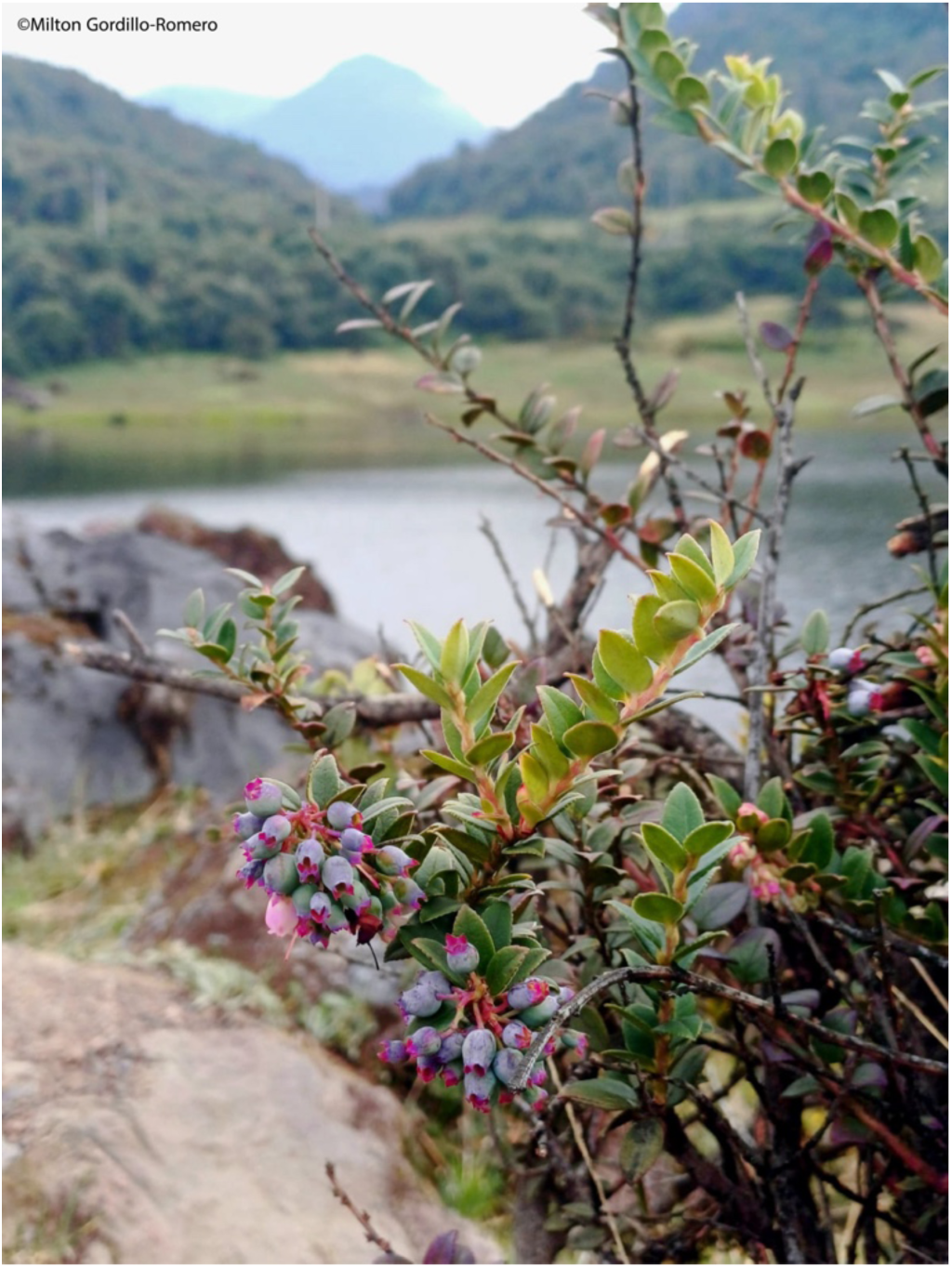
Plant of mortiño in the paramo of Papallacta, Ecuador

*V. floribundum* is a resilient species that has successfully adapted to high-altitude ecosystems, including the challenging conditions of the paramo in the Andes [16]. This distinctive mountain neotropical ecosystem of the northern Andean region is characterized by altitudes starting from 3,200 m.a.s.l to the lower limit of perpetual snow and glaciers [17]. It is characterized by non-arboreal vegetation and extreme environmental conditions such as freezing temperatures at higher elevations (>4,000 m.a.s.l) during the night, and intense ultraviolet radiation (UV index >11) during the day due to its geographical location flanking the equatorial line [18–21]. In this context, the Andean blueberry has evolved remarkable adaptations to withstand frost, freezing temperatures, and intense UV exposure, being one of the first plant species to regenerate following deforestation and human-induced fires in the paramo ecosystems [5,20,21].

Unlike the cultivated relatives of *Vaccinium*, there are not many studies of *V. floribundum* at the genetic and genomic level. However, a study carried out in the Ecuadorian highlands has revealed a high genetic diversity (He=0.73) and a clear population structure, including a unique cluster that groups Andean blueberry individuals growing above 4,000 m.a.s.l [6]. Furthermore, a recent study by [7] assembled the first chloroplast genome of *V. floribundum* and explored the phylogenetic relationships with other *Vaccinium* species, proposing *V. myrtillus* as the species most closely related to *V. floribundum*. With the constant development and optimization of sequencing technologies, obtaining draft genomes and chromosome-level assemblies for several commercially important and wild species within the genus *Vaccinium* has become increasingly more common. The currently published *Vaccinium* genomes: *V. macrocarpon* (American cranberry), *V. microcarpon* [22], *V. myrtillus* (bilberry) [23], *V. corymbosum* (blueberry) [24], *V. darrowii* [25], and *V. bracteatum* (Chinese wild blueberry) [26], have unraveled the phylogenetic relationships among the genus but have also shed light on the genetic mechanisms underpinning crucial agronomical traits, such as enhanced fruit quality, superior nutrient profiles and resilience to abiotic stressors [22,23,25,26]. However, most of these studies have focused primarily on *Vaccinium* species native to the northern hemisphere that grow in temperate climates. Obtaining the genome of *Vaccinium* species from other latitudes and ecosystems, such as the paramo, would provide valuable information to understand the evolutionary relationships and genetic mechanisms underlying adaptations within this genus. In this context, *V. floribundum*, adapted to paramo conditions, would be of special interest considering the global scenario of climate change.

In this research, we present the first high-quality reference genome of *V. floribundum*. The genome at the scaffold level was generated through a combination of nanopore-based long-reads and Illumina short reads. This approach yielded a final assembly with a genome size of 529 MB, containing 1,317 contigs and 97% complete BUSCOs, indicating a high level of completeness of the genome. The LTR Assembly Index (LAI) score of 12.93, reflects the quality of the assembled genome. The reference genome provided in this study constitutes a key tool for elucidating the genetic mechanisms underpinning the adaptations of *V. floribundum* to its life in the paramo ecosystem. Such insights are important for understanding this species’ ecological niche, role and evolution. This information could contribute to the conservation of this unique Andean endemic species; thereby safeguarding its ecological integrity and cultural significance.

## Methods

### Plant material and HMW-DNA extraction

Fresh young leaves were collected from a wild *V. floribundum* individual in the town of Lloa, Pichincha in the Ecuadorian Highlands (S0° 11.55122’ W78° 35.216’). Leaves were preserved in silica gel and transported to the Plant Biotechnology Lab at Universidad San Francisco de Quito (USFQ) and stored at −80 °C until DNA extraction. The sampling process was conducted under the Genetic Resource Permit Number: MAE-DNB-CM-2016-0046-M-0002 granted by the Ministerio del Ambiente, Agua y Transición Ecológica in Ecuador, per Ecuadorian law. The sampled individual was formally registered in the USFQ Herbarium (QUSF) with the accession code 079. The CTAB-Genomic Tip (Oxford Nanopore Technologies) protocol was used to isolate High Molecular Weight DNA (HMW-DNA) from 2g of young leaves. Subsequently, the DNA was purified using the QIAGEN Genomic-tip 100/G affinity columns and resuspended in 100 μl of TE buffer. Finally, HMW-DNA enrichment was performed using the short-read eliminator kit SRE (Circulomics) to deplete DNA fragments shorter than 25Kb.

### Oxford Nanopore Sequencing

Six genomic libraries were prepared according to the Ligation Sequencing Kit SQK-LSK109 (Oxford Nanopore Technologies) workflow, using 2 μg of DNA as initial input. Purification of the libraries after each step was carried out with AMPure XP beads (Beckman) and quantified by fluorometry using Qubit 3.0 (Thermo Fisher Scientific). The final libraries with 50 fmol of DNA were sequenced for 24 hours on 3 flowcells (R9.4.1) in a MinION Mk1b sequencer with the fast base-calling algorithm executed with Guppy v5.1.13 (Oxford Nanopore Technologies).

The resulting fastq files from each sequencing run were compiled in a single file and sequencing adapters were removed using Porechop v0.2.4 (RRID:SCR_016967) [27]. Reads with quality scores <7 were filtered out using Nanofilt v2.8.0 (RRID:SCR_016966) [28]. The final raw read dataset statistics were evaluated using LongQC v1.2.0c [29] and Nanoplot v1.33.0 (RRID:SCR_024128) [28].

### Illumina Sequencing

To complement the long-read sequencing dataset, short-read Illumina sequencing was performed. Two μg of genomic DNA were sent to BGI (Hong Kong, China) for NovaSeq 150PE sequencing. Quality check of the data was performed using FastQC (RRID:SCR_014583) [30].

### Genome size estimation

The genome size and heterozygosity for *V. floribundum* were estimated using a k-mer-based analysis (21-mers) with Jellyfish v2.3.0 (RRID:SCR_005491) [31]. Trimmed and filtered reads from Illumina DNA libraries were employed as input. The resulting k-mer profiles were visualized with GenomeScope v2.0 (RRID:SCR_017014) [32].

### *De novo* genome assembly

*De novo* assembly of ONT reads was performed using SMARTdenovo v1.0.0 (RRID:SCR_017622) [33] and Flye v2.9.2 (RRID:SCR_017016) [34] with default parameters. The assemblies were first polished using ONT raw reads with one round of Medaka v1.11.1 (Oxford Nanopore Technologies) and another round using Illumina short reads with POLCA, the built-in polisher of the Maryland Super-Read Celera Assembler v.4.1.0 (MaSuRCA, RRID:SCR_010691) [35]. Additionally, a hybrid assembly was generated with MaSuRCA v.4.1.0 (RRID:SCR_010691); a hybrid *de novo* assembler that constructs the assembly using short reads and polishes and scaffolds the contigs using long reads [35], using default parameters. This assembly was also polished using ONT raw reads with one round of Medaka v1.11.1 (Oxford Nanopore Technologies) and another round using Illumina short reads with POLCA (MaSuRCA, v4.1.0 RRID:SCR_010691) [35].

### Genome assembly quality, continuity and completeness assessment

Comparisons of quantitative and qualitative metrics corresponding to the obtained assemblies were performed to evaluate their quality, continuity and completeness with Quast v5.2.0 (RRID:SCR_001228) [36], using *V. myrtillus* (GCA_016920895.1) as reference and BUSCO v5.4.7 (RRID:SCR_015008) using the eudicots_odb10 database which comprises 2,326 total genes [37]. *V. myrtillus* (GCA_016920895.1) was used as reference due to its close phylogenetic relationship to *V. floribundum* [38].

The Long Terminal Repeat (LTR) Assembly Index (LAI) was also used to evaluate assembly continuity and completeness of non-sequencing, repeat regions of the genome [39]. LTRharvest v1.6.2 (RRID:SCR_018970) [40] and LTR_FINDER v1.0.7 (RRID:SCR_015247) [41] were first employed to identify intact LTR retrotransposons and create an LTR sequencing library. LTRharvest was run with the following parameters: -minlenltr 100 -maxlenltr 7000 -mintsd 4 - maxtsd 6 -motif TGCA -motifmis 1 -similar 85 -vic 10 -seed 20 -seqids yes. The LTR sequence library was then used in LTR_retriever v2.8.7 (RRID:SCR_017623) [42] with default parameters to calculate the LAI scores of the assemblies.

Additionally, to examine the uniformity and distribution of the coverage of ONT and Illumina reads along the *V. floribundum* genome, the longest contig of each assembly was extracted using the *samtools faidx* function of the SAMtools package v1.18 (RRID:SCR_002105) [43]. The longest contig was then mapped to the ONT reads using minimap2 v2.26 (RRID:SCR_018550) [44] and to the Illumina reads using BWA (RRID:SCR_010910) [45] under default parameters. To obtain the coverage of reads in each position, the *samtools depth* function of the SAMtools package v1.18 (RRID:SCR_002105) [43] was employed. The graph was then plotted using a previously described script [38].

### Genome assembly selection, read filtering and genome scaffolding

The best assembly was selected based on the metrics obtained from the genome assembly quality, continuity and completeness assessment. To further improve this assembly and to reduce possible contaminations, ONT raw reads of less than 1,000 bp were filtered out before following the same assembly and polishing pipeline steps described in the “*De novo* genome assembly” section. The polished assembly was then scaffolded using ntLink v1.3.9 [46] with the gap_fill option and the overlap=true parameter. After scaffolding, the assembly was screened for foreign contaminations using the NCBI Foreign Contamination Screen tool (FCS_GX) [47], and the contaminant sequences were removed with the same tool.

The final resulting assembly was then assessed for quality, continuity and completeness using the tools described in the “Genome assembly quality, continuity and completeness assessment” section.

### Genome annotation

The final resulting assembly was annotated. RepeatModeler v2.0.3 (RRID:SCR_015027) [48] was first used to construct a *de novo* custom repeat library of the assembly. The “LTRStruct” option for long terminal repeat (LTR) retroelement identification was specified. Maker v3.01.03 (RRID:SCR_005309) [49] was then used for genome annotation. Repetitive regions were identified and soft-masked with RepeatMasker v4.0.9_p2 (RRID:SCR_012954) [50]. The masked genome was then annotated in three consecutive rounds. *Ab initio* gene prediction algorithms were run with EST and protein evidence in the first round using the est2genome and protein2genome parameters. Protein sequences from *Vaccinium darrowii* [51] and EST data obtained from the NCBI EST database (2023) for all available *Vaccinium* species were used as evidence. In the second round of annotation, the *ab initio* gene predictor SNAP [52] was trained with the initial predictions of round one before running Maker employing the hidden Markov model (HMM) from SNAP. Finally, a third round of annotation was run with SNAP. All three consecutive rounds of Maker generated protein and transcript fasta files and gff files that were merged before identifying the best-supported gene models. To isolate the best-supported gene models, InterProScan v5.64-9.60 (RRID:SCR_005829) [53] was run to identify conserved Pfam domains on the Maker-predicted proteins. Using accessory scripts from Maker, gene models with AED values lower than 1 or lacking Pfam domains were removed from the gff and fasta files. Finally, to obtain the annotation statistics, the agat_sp_statistics.pl script from AGAT (Another Gff Analysis Toolkit) v1.2.0 [54] was run.

## Results & Discussion

### DNA sequencing results

Nanopore sequencing assays generated a total of 12.99 Gb corresponding to 3,623,409 reads, equivalent to approximately ∼25x coverage of the *V. floribundum* genome (∼520 Gb, based on a closely related *Vaccinium* species) [23]. In general, reads had a mean quality score of 8.9 and a mean read length of 3.58 Kb. In addition, NovaSeq Illumina sequencing assays generated a total of 96.62 Gb equivalent to 644,138,748 reads with a quality score >q20. This is equivalent to ∼186x coverage of 150-bp paired-end reads. The combined long and short read data obtained in the sequencing step provided sufficient coverage for assembling the *V. floribundum* genome.

### Genome size and heterozygosity estimation

Using the Illumina short reads, the genome size estimate for *V. floribundum* was calculated using a k-mer analysis with a k-value of 21. The analysis estimated a genome size of 408 Mb (**Additional File 1 - Figure S1**), whereas the final genome assembly was 529 Mb. This final genome size is comparable with other available *Vaccinium* species (*V. darrowii*: 522 Mb [25]; *V. corymbosum:* 393 Mb [55]; *V. macrocarpum:* 490.68 Mb [22]). The *V. floribundum* genome had a single k-mer frequency peak consistent with a diploid genome [56]. Additionally, its estimated heterozygosity (0.223%) was relatively low when compared to the reported values for *V. darrowii* (1.27%) [25] and *V. macrocarpon* (0.863 – 0.934%) [22]. This could be related to the findings of a study that reported that wild populations of cranberry (a *Vaccinium* species) exhibited lower levels of heterozygosity than cultivated ones [57]. The same phenomenon could be occurring in this case, as *V. floribundum* is a wild species of high-altitude ecosystems. However, other factors like sequencing depth and repeat content could also be contributing to the differences in heterozygosity values [58].

### *De novo* genome assembly quality and accuracy assessments

From the initial assemblies carried out and polished with both ONT and Illumina reads, both the SMARTdenovo and MaSuRCA assemblies were similar in size (509 Mb and 511 Mb) while the Flye assembly was slightly larger with a total genome length of 533 Mb. The largest contig in the Flye assembly was 9.5 Mb with an average N50 size of 2.1 Mb. These values were significantly higher than those found in the other two assemblies where the largest contig in the SMARTdenovo assembly was 2.7 Mb and the largest contig in the MaSuRCA assembly was 5.7 Mb. As for the N50, the SMARTdenovo assembly had an N50 average size of 633 kb while the MaSuRCA assembly had an average N50 of 788 kb. Despite the Flye assembly presenting better genome parameters than the other assemblies, this assembly also generated around twice the number of contigs (3,865 contigs) than the SMARTdenovo (1,556 contigs) and MaSuRCA (1,180 contigs) assemblies. Nonetheless, the Flye assembly presented the lowest L50 value of the three generated assemblies (82 contigs compared to 246 contigs for the SMARTdenovo assembly and 174 contigs for the MaSuRCA assembly) (**Table 1**).

**Table 1.**
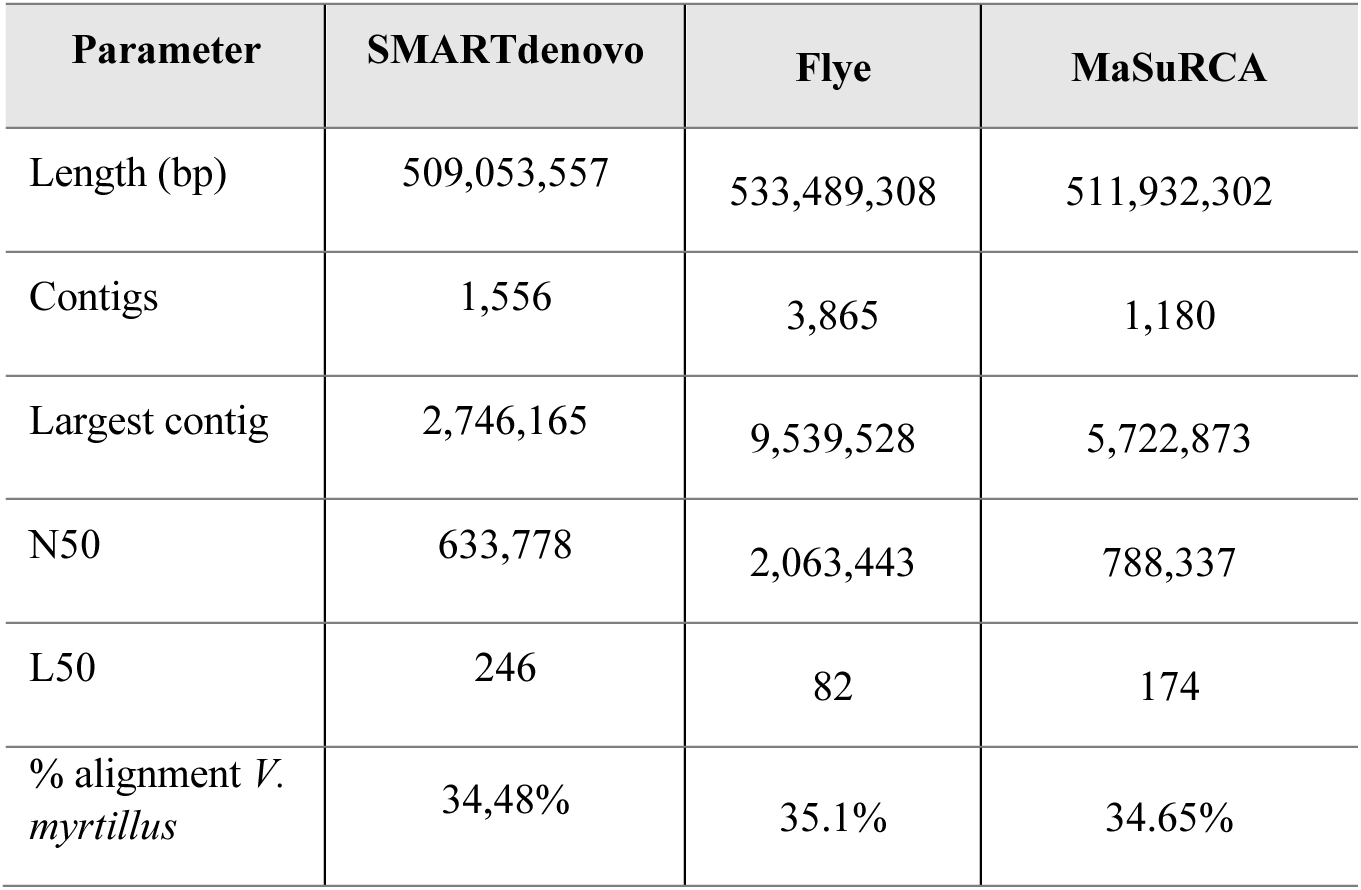
Quantitative measurement assessments between initial assemblies of *V. floribundum*.

To further compare the three generated assemblies, the three draft genomes were aligned to the reference genome of a closely related species of the same genus, *V. myrtillus* (GCA_016920895.1). The percentage alignment to the reference genome was similar for all three assemblies with the Flye assembly being slightly higher (35.1%) (**Table 1**).

The Long Terminal Repeat (LTR) Assembly Index (LAI) metric was also used to evaluate the continuity and genome assembly quality of non-coding, repeat regions. Higher LAI scores usually correspond to more continuous and complete assemblies since they often contain a larger number of intact LTR retrotransposons [42]. The SMARTdenovo assembly had an LAI score of 9.70, which characterizes it as a draft genome. In comparison, the MaSuRCA (11.68) and Flye (14.10) assemblies had higher LAI Scores. Genomes with LAI scores between 10 and 20 are considered genomes of reference quality [42] (**Table 2**).

**Table 2.** LAI score indexes and genome assembly categories of *V. floribundum*.

| Assembly | LAI Score | Category [42] |
| --- | --- | --- |
| SMARTdenovo | 9.70 | Draft |
| Flye | 14.10 | Reference |
| MaSuRCA | 11.68 | Reference |

We also used BUSCO to assess the completeness of the genome. To do so, BUSCO genes from the eudicot_odb10 database (2,326 genes) were analyzed in the three assemblies. The analysis showed similar trends in the three assemblies. The Flye assembly had the highest percentage of Complete (C) BUSCO genes (96.9%). Nonetheless, the Complete (C) BUSCO genes for the SMARTdenovo and MaSuRCA assemblies were only slightly lower with 95.8% and 96.1% complete genes, respectively (**Figure 2**).

**Figure 2.**
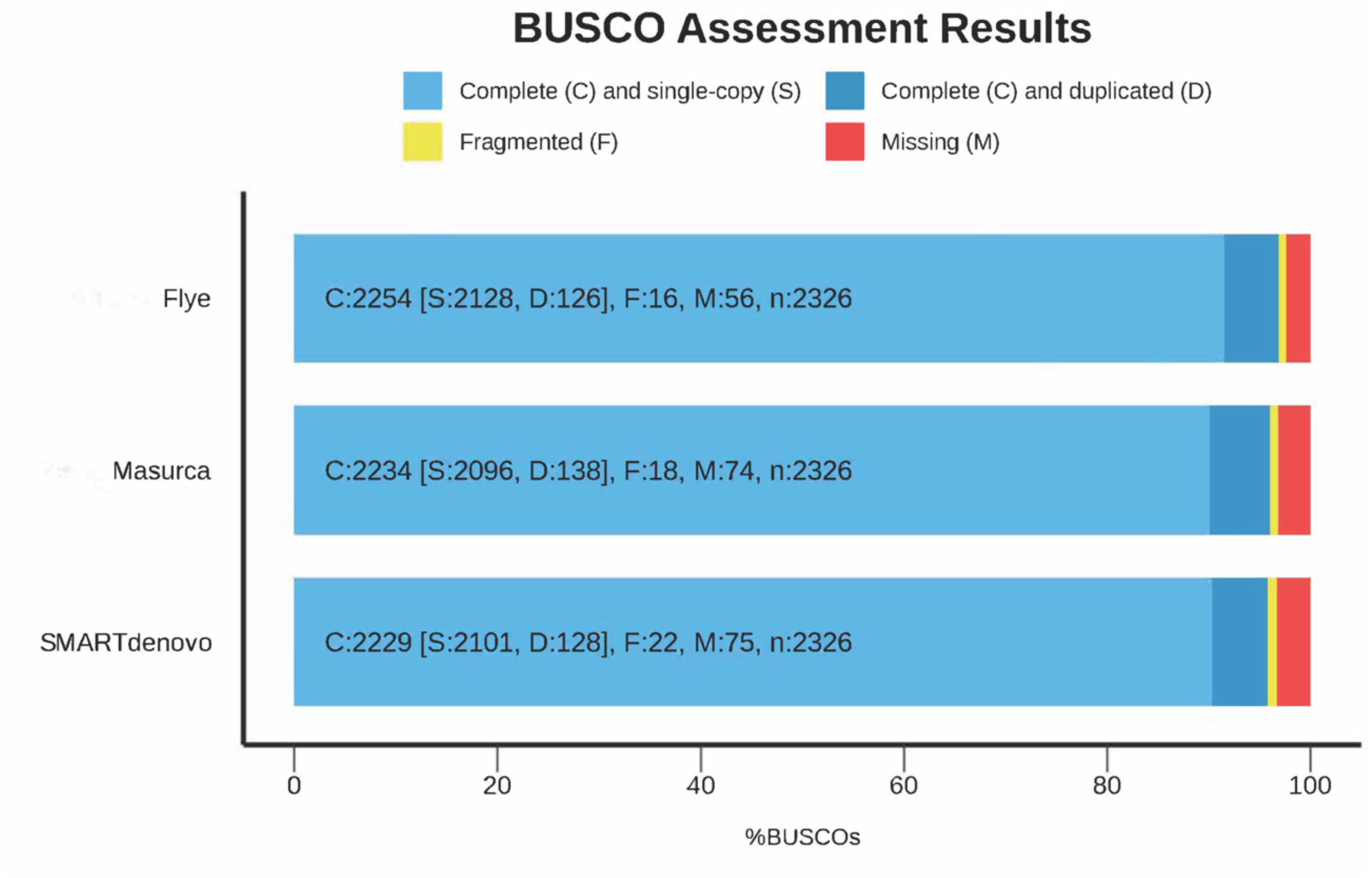
BUSCO assessment results. Comparisons between the Flye, MaSuRCA, and SMARTdenovo assemblies using de eudicots_odb10 database (n = 2326).

When the ONT and Illumina reads were mapped independently to the longest contig of the three assemblies, the obtained coverage graphs showed similar results between the three, indicating suitable *de novo* assembly results. Interestingly, the coverage graphs generated with the Illumina reads showed a more consistent distribution of reads along the analyzed region of the genome than the graphs obtained with the ONT reads. Additionally, the even distribution of reads in the Flye assembly was notably better than in the other generated assemblies (**Additional File 1 - Figure S2; Additional File 1 - Figure S3**).

Flye consistently outperforms other long-read assembly tools such as CANU, Miniasm, Raven and Redbean [59]. Particularly for plant genomes, the Flye and CANU assemblers are usually recommended for achieving accurate and complete assemblies [60]. Flye’s algorithm employs a uurepeat graph construction approach that is adept for handling high error rates and complex repeat regions typical of long-read sequencing data, enabling it to produce more accurate and contiguous assemblies [61]. In contrast, SMARTdenovo and MaSuRCa are more general-purpose assemblers that are not specifically tailored for long-read data. While they can handle a combination of long and short reads, their algorithms may not be as effective at resolving the complexities inherent in long-read datasets, particularly when dealing with highly repetitive plant genomes [33,62].

### Establishing and evaluating the best assembly

Based on the analyzed metrics, the Flye assembly was selected as having the best quality genome and was chosen for further processing and analyses. After filtering out the ONT reads shorter than 1,000 bp, screening and eliminating foreign contaminants, assembling and scaffolding, the final reference genome was obtained (**Figure 3**).

**Figure 3.**
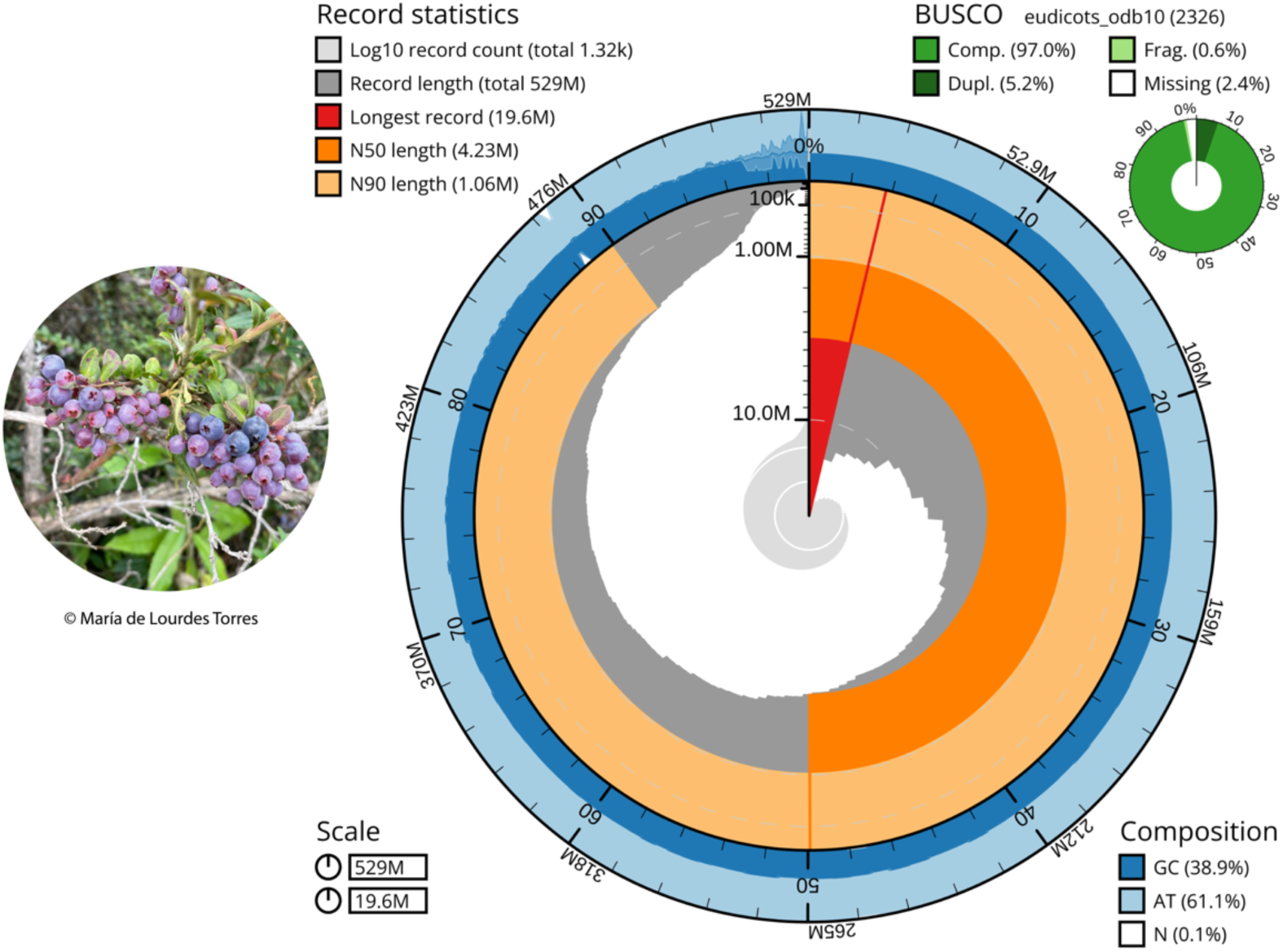
Snail Plot (Blobtoolkit) describing and representing the assembly statistics of the final genome assembly of *V. floribundum*. The plot shows N50, N90, longest contig and total genome length metrics as well as BUSCO scores obtained from the eudicots_odb10 database. GC% content is displayed in the outer ring.

This assembly contains 1,317 contigs, fewer contigs than the previous draft assembly and comparable to other *Vaccinium* genome assemblies, including chromosome-level assemblies [22,23,25,56]. The total genome length of our final assembly is 529 Mb, which falls within the range of genome sizes found in *Vaccinium* species [22,23,25,55,56] as seen in the Genome Size Estimation section. Furthermore, the obtained genome size is very similar to that of its closest relative within the genus, *V. myrtillus*, which has an estimated genome size of 524.3 Mb [7,23]. Our final *V. floribundum* assembly improved in several statistics following read filtering. For example, its largest contig size is 19.6 Mb and has an average N50 size of 4.2 Mb, higher than some reported *Vaccinium* genomes (*V. oxycoccos*: 1.8 Mb [56]; *V. microcarpum*: 176.33 kb [22]). Additionally, the assembly’s L50 has been reduced to 37 contigs, and its L90 is represented in only 133 contigs (**Figure 3**).

To further validate the results, the LAI score index was also calculated. The final assembly had an LAI score of 12.93, classifying it as a reference genome [42] and being comparable to other reference *Vaccinium* genomes that report LAI scores from 13.3 (*V. corymbosum*) [55] to 17.57 (*V. macrocarpum*) [22]. The BUSCO assessment was also performed for this final assembly; however, the results did not vary (complete BUSCOs: 97%) (**Figure 3**) compared to those obtained before read filtering, contamination removal and genome scaffolding. The BUSCO values obtained in the final assembly are an indicator of a continuous genome and are slightly higher to most reported *Vaccinium* genome assemblies [22,25,56].

ONT and Illumina reads were also mapped to the longest contig of the final assembly to evaluate the evenness of reads along the analyzed region of the genome (**Figure 4**). The obtained coverage graphs showed a more consistent distribution of reads, both with ONT and Illumina reads than what was obtained in the previous version of the Flye assembly (**Additional File 1 - Figure S2; Additional File 1 - Figure S3**).

**Figure 4.**
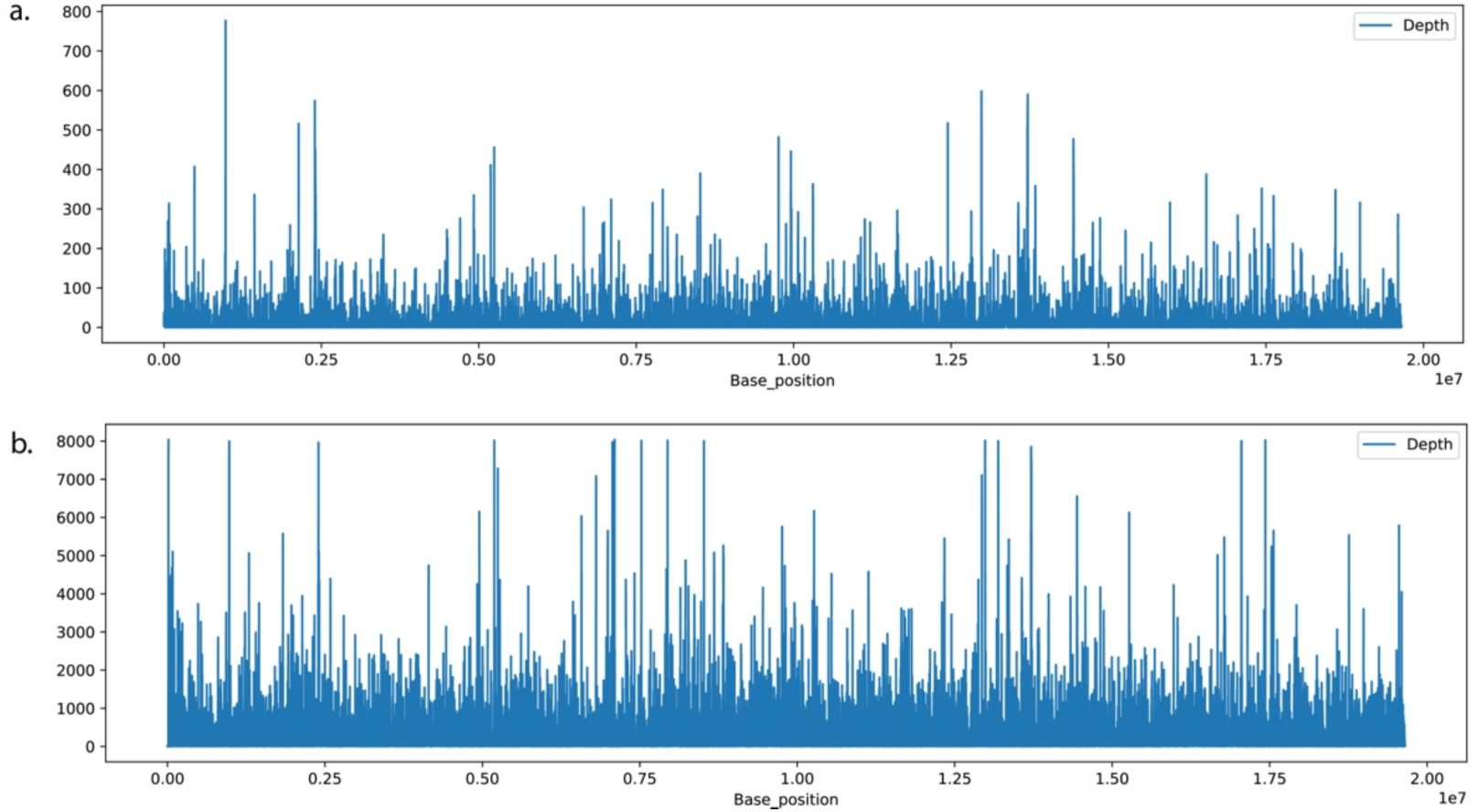
Depth coverage graph of the final assembly of *V. floribundum* of the largest contig using **a.** ONT raw reads and **b.** Illumina raw reads.

All together the metrics and parameters obtained of the *V. floribundum* genome assembly allow us to report it as a high-quality continuous genome comparable to other published *Vaccinium* genome assemblies [22,25,56].

### Genome Annotation

The annotation of the final genome of *V. floribundum* in Maker predicted a total of 33,847 protein-coding genes when eliminating predicted gene models with AED values greater than 1 (**Table 4**). AED values closer to 1 usually represent little to no agreement between the annotated gene models and the protein/ transcript evidence while values closer to 0 show greater consensus [63].

**Table 4.** Summary statistics of the annotated final genome of *V. floribundum*.

| Statistic | *V. floribundum* final genome |
| --- | --- |
| Number of genes | 33,847 |
| Number of exons | 167,341 |
| Number of introns (in CDS) | 133,056 |
| Overlapping genes | 1940 |
| Mean mRNAs per gene | 1.0 |
| Mean exons per mRNA | 4.9 |
| Mean introns per mRNA | 3.9 |
| Mean gene length (bp) | 6,574 |
| Mean exon length (bp) | 220 |
| Mean intron length (bp) | 1,391 |

The resulting 33,847 protein-coding genes of the *V. floribundum* genome are comparable to what has been reported for other *Vaccinium* species (*V. darrowii*: 34,809 [51], *V. microcarpum*: 30,147 [22]; *V. corymbosum:* 33,183 [55]. Interestingly, the closest *Vaccinium* species to *V. floribundum*, *V. myrtillus* [7], presented not only a similar number of protein-coding genes (36,404), but also a mean exon length of 209 bp [23], very similar to our obtained mean exon length (220 bp) for *V. floribundum*. Nonetheless, it is interesting to note that reported protein-coding genes are variable within other *Vaccinium* species going from 23,532 genes in *V. macrocarpon* [22] to *V. oxycoccos* with 50,621 genes [56].

Additionally, the annotation of the final genome of *V. floribundum* presented an average gene length of 6,574 bp, similar to what has been reported for *V. darrowii* (5,076.09 bp) [25] and *V. corymbosum* (4,554 bp) [55]. Similarly, the number of exons per gene (4.9) is consistent with what has been reported in other *Vaccinium* species (*V. darrowii*: 5.02 [25]; *V. macrocarpom*: 6.7 [22]; *V. microcarpum*: 6.9 [22]; *V. corymbosum*: 5.3 [55]) as well as the average exon length (220 bp) (*V. corymbosum*: 238 bp [55], demonstrating a good and reasonable quality of the annotation.

### Conclusions

In this study, we report the first reference genome for *V. floribundum*, an endemic and resilient species from the Andean region. This high-quality genome was sequenced and assembled using Oxford Nanopore long reads and Illumina short reads. This novel genomic resource is comparable in terms of size, completeness and quality to other reference genomes already reported for other *Vaccinium* species, thus contributing to the current knowledge within the genus. The reference genome of this wild species adapted to unique environmental conditions of the Andean tropics represents a relevant molecular resource that will facilitate comprehensive research on the phylogenetic and evolutionary relationships of *V. floribundum.* Finally, it constitutes the basis for future genomic studies aiming to understand Andean blueberry adaptations to extreme altitude ecosystems, but also for genomic population studies essential to the conservation of this emblematic species.

## Additional Files

### Additional File 1 (.pdf)

**Figure S1:** k-mer analysis used to estimate *V. floribundum* genome size and heterozygosity and visualized in GenomeScope. K-mer size was set at 21.

**Figure S2:** Depth coverage graph of *V. floribundum* of the largest contig using ONT raw reads of **a.** SMARTdenovo assembly, **b.** MaSuRCA assembly, and **c.** Flye assembly.

**Figure S3:** Depth coverage graph of *V. floribundum* of the largest contig using Illumina raw reads of a. SMARTdenovo assembly, b. MaSuRCA assembly, and c. Flye assembly.

## Data Availability

The genome assembly and all sequencing data have been deposited in the GenBank database under the BioProject PRJNA1071645 (SAMN39706721). The script used for the assembly and annotation of this genome is described in protocol.io (dx.doi.org/10.17504/protocols.io.n92ldmo4nl5b/v1). All supporting data and material is available in the *GigaScience* GigaDB database.

## Abbreviations

m.a.s.l: meters above sea level
UV: ultraviolet light
BUSCO: Benchmarking Universal Single-Copy Orthologs
LAI: Long Terminal Repeat Assembly Index
QUSF: USFQ herbarium
HMW-DNA: High Molecular DNA
CTAB: Cetyltrimethylammonium bromide
SRE: short-read eliminator
ONT: Oxford Nanopore Technologies
MaSuRCA: Maryland Super-Read Celera Assembler
QUAST: Quality Assessment Tool for Genome Assemblies
LTR: long terminal repeat
BWA: Burrows-Wheeler Aligner
NCBI: National Center for Biotechnology Information
EST: Expressed sequence tag
HMM: hidden Markov model
gff: general feature format
AGAT: Another Gff Analysis Toolkit
Gb: gigabase
Kb: kilobase
bp: base pair
Mb: megabase
AED: Annotation Edit Distance
CDS: coding sequence

## Author’s Contributions

MLT conceived and designed the study. MAQ, GP, MGR and CEA performed the experiments, analyzed, and interpreted the assemblies and annotations. MAQ, GP, MGR and CEA wrote the draft of the manuscript and MLT revised, edited and improved the manuscript. All authors contributed to and approved the final manuscript.

## Acknowledgments

The authors are grateful to Estefanía Rojas, Sebastián Jordán and Manuela Parra for their contributions to the overall project. A special thanks to Colegio de Ciencias Biológicas y Ambientales from Universidad San Francisco de Quito that without their funding we would not have been able to carry out this study.

