## Additional File 1 for "First whole genome assembly of *Vaccinium floribundum* Kunth, an emblematic Andean species"

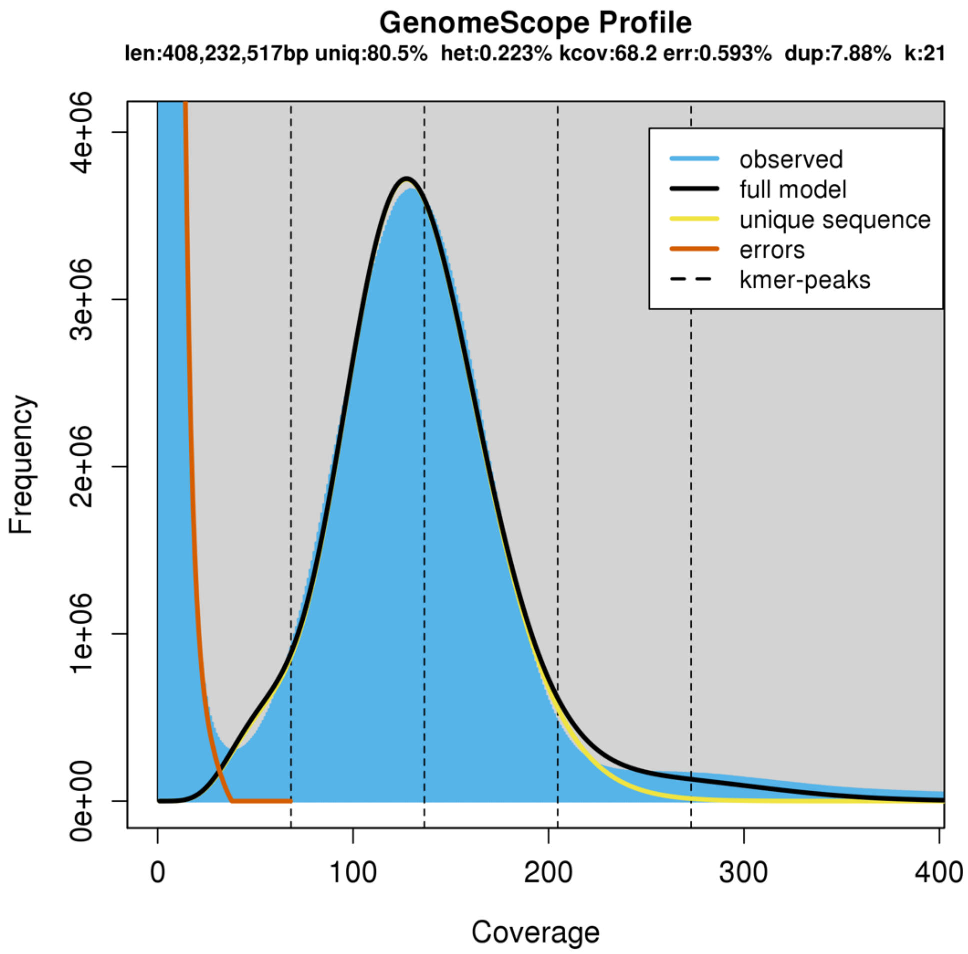


**Figure S1.** k-mer analysis used to estimate *V. floribundum* genome size and heterozygosity and visualized in GenomeScope. K-mer size was set at 21.

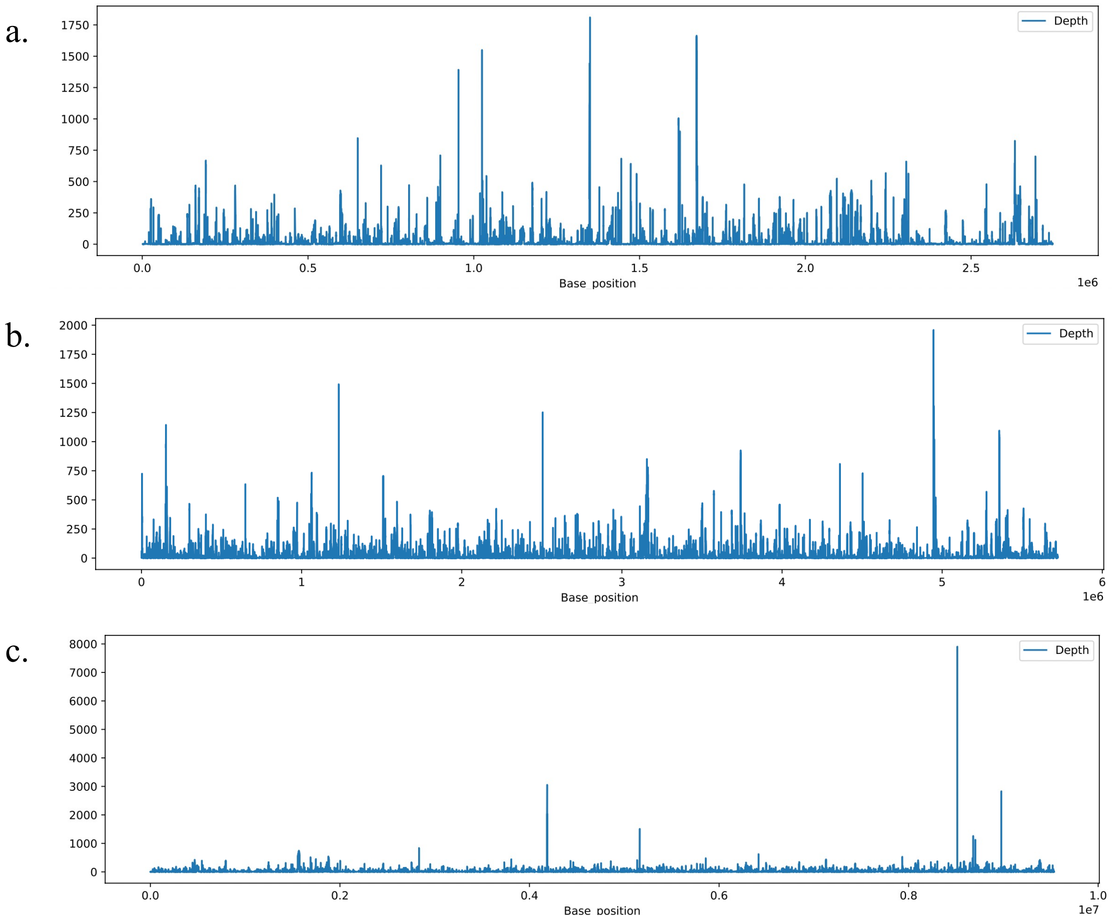


**Figure S2.** Depth coverage graph of *V. floribundum* of the largest contig using ONT raw reads of **a.** SMARTdenovo assembly, **b.** MaSuRCA assembly, and **c.** Flye assembly.

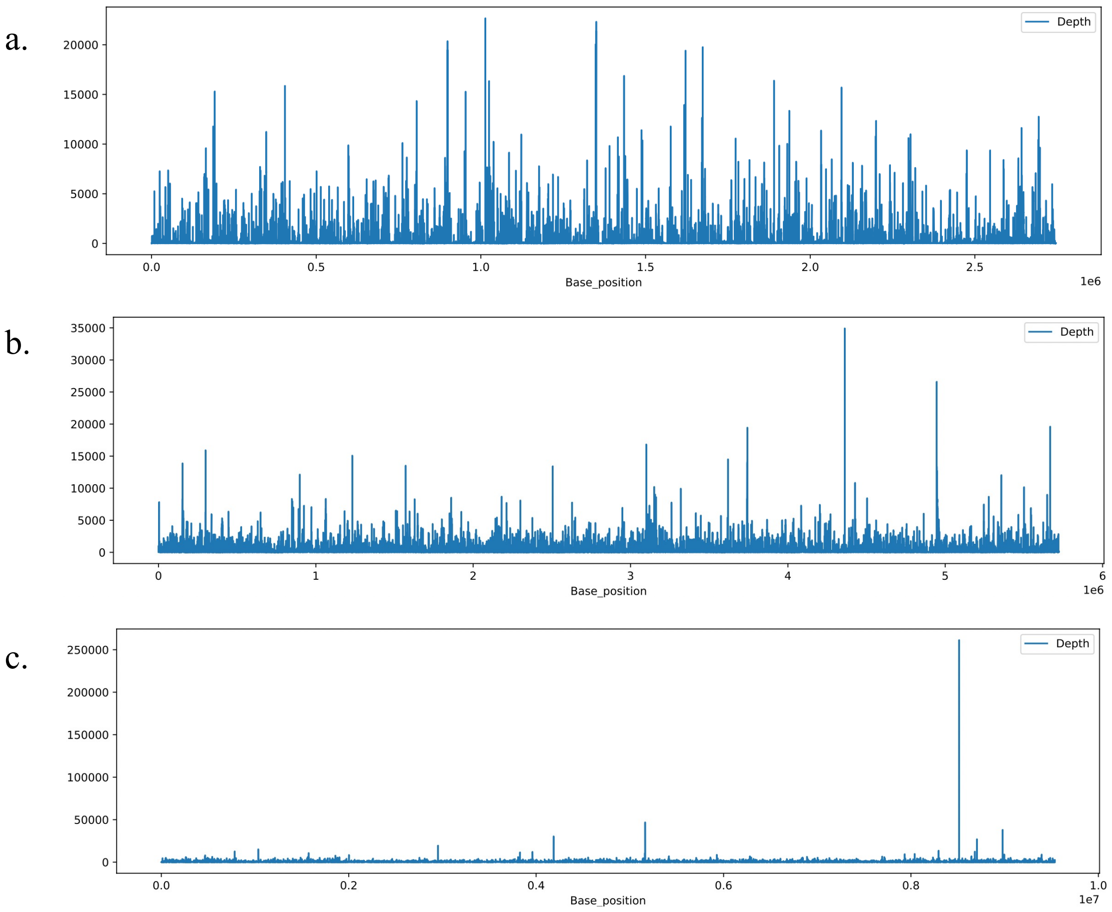


**Figure S3.** Depth coverage graph of *V. floribundum* of the largest contig using Illumina raw reads of **a.** SMARTdenovo assembly, **b.** MaSuRCA assembly, and **c.** Flye assembly.
